# Quantification of iPSC-derived vascular networks in novel phototunable angiogenic hydrogels

**DOI:** 10.1101/2020.08.25.259630

**Authors:** Cody O. Crosby, Alex Hillsley, Sachin Kumar, Sapun H. Parekh, Adrianne Rosales, Janet Zoldan

## Abstract

Vascularization of engineered scaffolds remains a critical obstacle hindering the translation of tissue engineering from the bench to the clinic. Previously, we demonstrated the robust micro-vascularization of collagen hydrogels with induced pluripotent stem cell (iPSC)-derived endothelial progenitors; however, physically cross-linked collagen hydrogels compact rapidly and exhibit limited strength. To address these challenges, we synthesized an interpenetrating polymer network (IPN) hydrogel comprised of collagen and norbornene-modified hyaluronic acid (NorHA). This dual-network hydrogel combines the natural cues presented by collagen’s binding sites and extracellular matrix (ECM)-mimicking fibrous architecture with the *in situ* modularity and chemical cross-linking of NorHA. We modulated the stiffness and degradability of this novel IPN hydrogel by varying the concentration and sequence, respectively, of the NorHA peptide cross-linker. Rheological characterization of the photo-mediated gelation process revealed that the stiffness of the IPN hydrogel increased with cross-linker concentration and was decoupled from the bulk NorHA content. Conversely, the swelling of the IPN hydrogel decreased linearly with increasing cross-linker concentration. Collagen microarchitecture remained relatively unchanged across cross-linking conditions, although the mere addition of NorHA delayed collagen fibrillogenesis. Upon iPSC-derived endothelial progenitor encapsulation, robust, lumenized microvascular networks developed in IPN hydrogels over two weeks. Subsequent computational analysis showed that an initial rise in stiffness increased the number of branch points and vessels, but vascular growth was suppressed in high stiffness IPN hydrogels. These results suggest that an IPN hydrogel consisting of collagen and NorHA is highly tunable, compaction resistant, and capable of stimulating angiogenesis.

**STATEMENT OF SIGNIFICANCE:** We have synthesized the first tunable collagen and norbornene functionalized hyaluronic acid (NorHA) interpenetrating polymer network hydrogel. This unique biomaterial allows for control over hydrogel stiffness, independent of the total polymer concentration, by varying the concentration of a peptide cross-linker and was specifically designed to produce a biomimetic vasculogenic microenvironment. Using the system, we performed a detailed study of the vasculogenesis of induced pluripotent stem cell-derived (iPSC) endothelial progenitors, a poorly studied cell source with considerable therapeutic potential. Our results show that vascular growth can be tuned by altering the stiffness and degradability of the scaffolds independently. Finally, we improved upon our open-source computational pipeline programmed in ImageJ and MATLAB to further quantify vascular topologies in three dimensions.

**Graphical abstract:** 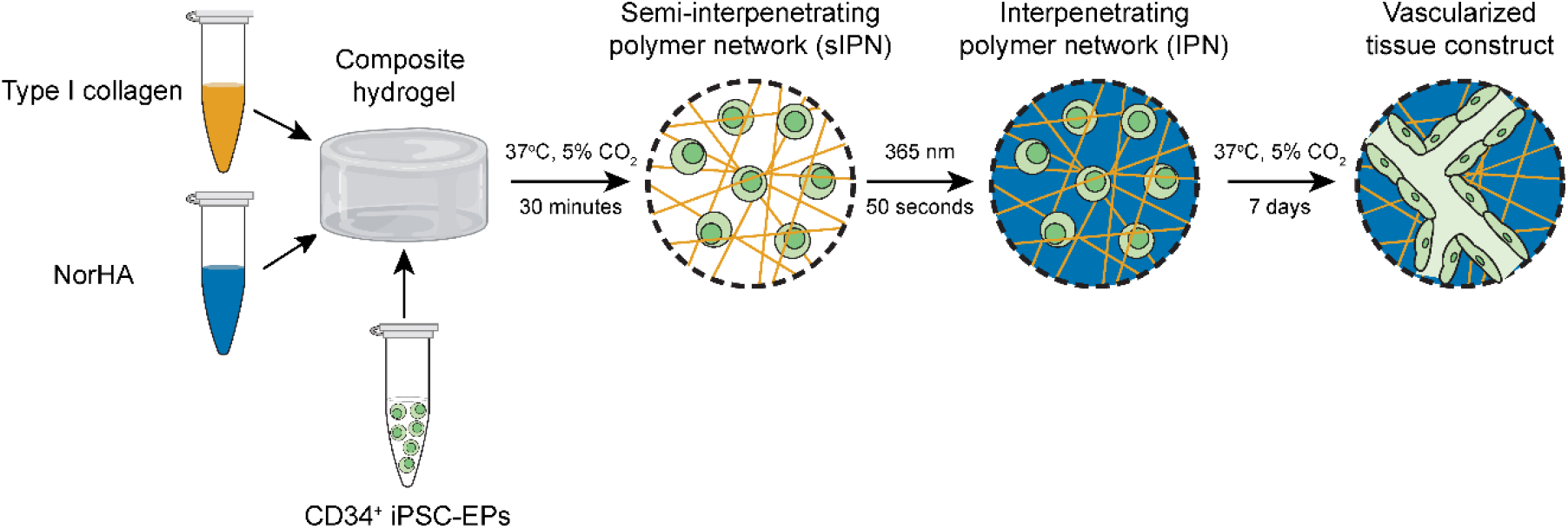

## 1. INTRODUCTION

The revascularization of lab-grown and native ischemic tissue remains one of the most critical challenges facing the fields of tissue engineering and regenerative medicine [1]. A practical *in vitro* model of vascular development promises to inspire new vascular therapies by enhancing our fundamental understanding of vasculogenic/angiogenic mechanisms and by constructing more accurate disease models for therapeutic drug testing [2,3]. Angiogenic biomaterials promise to be vital components of revolutionary platforms, such as bio-printing and organ-on-a-chip, that may uncover molecular and tissue-level mechanisms of cardiovascular disease [4,5]. Biomaterial-based platforms that seek to mimic the vascular milieu typically incorporate primary endothelial and perivascular cells to recapitulate physiological microvasculature [6]. Since these cells are often sourced from different patients and animal models, comparing the generated microvasculature across published studies is difficult; therefore, the reproducibility of such studies is often severely compromised.

In contrast, endothelial progenitors (EPs) are bipotent and can, therefore, be further differentiated into endothelial cells and smooth muscle cells/pericytes, the building blocks of mature, functional vasculature [7]. However, the critically low number of functional EPs present in tissues such as bone marrow is therapeutically limiting. Also, patients with cardiovascular disease (CVD) and diabetes often possess EPs with reduced functionality [8–11]. In contrast, induced pluripotent stem cells (iPSCs) are a patient-specific, largely untapped cell source that can generate functional somatic cells from patients with risk factors, as was shown for iPSCs derived from diabetic patients [12]. Additionally, iPSCs have recently emerged as patient-specific therapies for many vascular conditions such as progressive diabetic retinopathy [13]. Different protocols to differentiate iPSCs into EPs have emerged in recent years, and EP differentiation efficacy has steadily risen as these protocols are progressively refined [14–16]. Despite these advances, the response of iPSC-EPs to variations of physical and chemical characteristics in the local extracellular matrix (ECM) remains relatively unknown and is a significant barrier to translation.

To this end, biomaterials provide an excellent means to study this process *in vitro*, due to their high tunability and ability to mimic native tissue [17]. We have previously utilized rat tail collagen to demonstrate the high vasculogenic potential of iPSC-EPs [18]. Collagen hydrogels are a convenient angiogenic biomaterial because collagen contains essential cell attachment motifs, is matrix metalloprotease (MMP)-sensitive, and naturally maintains a micro-scale fibril architecture [19]. However, it remains challenging to decouple stiffness, fiber density, and degradability in collagen hydrogels, as these parameters are all dependent on the bulk polymer density. Additionally, since these collagen hydrogels are physically cross-linked, they readily compact with exposure to increasing cell contractility and MMP-mediated matrix remodeling, which limits their culture time and use in longitudinal studies. The ability to decouple stiffness and fiber density would allow for the creation of a mechanically tunable 3D scaffold that could unravel new mechano-instructive drivers of vasculogenesis [20]. We chose to use NorHA as the co-polymer in our system because NorHA’s stiffness, degradability, and bioactivity can be readily tuned independently of its concentration.

Hyaluronic acid (HA) is a naturally occurring biopolymer that can be functionalized and cross-linked to form hydrogels [21] with a wide range of tunable properties, including stiffness [22], signaling gradients [23], degradability [24], and viscoelasticity [25]. Of interest in this study, hyaluronic acid functionalized with norbornene groups (NorHA) has been demonstrated to allow for independent control over stiffness and degradation [26]. Utilizing thiol-ene click chemistry, NorHA is cross-linked via the incorporation of a di-thiol cross-linker, such as a peptide with two cysteine residues. Recent work has highlighted precise control over peptide degradation rates by varying the amino acid sequence of this di-thiol cross-linker [27]. By incorporating variably degradable peptide cross-linkers, one can translate this control of peptide degradability to control over hydrogel degradability [28]. However, HA hydrogels (including NorHA) lack the macroscale fibrillar structure necessary for significant network development [29].

To probe the effects of varying stiffness and degradability on iPSC-EP network development, we describe the synthesis of a novel interpenetrating polymer network (IPN) of NorHA and type I collagen. IPN hydrogels of independently cross-linked methacrylated HA and collagen have been synthesized for 3D 3T3 fibroblast cell culture [30], 2D Schwann cell culture [31], and pancreatic islet transplantation [32]. In a recent groundbreaking study, Lou *et al.* utilized an HA-hydrazone and collagen IPN to demonstrate the effects of viscoelasticity on mesenchymal stem cell (MSC) spreading and focal adhesion formation [33]. To our knowledge, vascular networks have not been cultured in HA-collagen IPN hydrogels; additionally, no previous study has studied the impact of HA incorporation on collagen architecture.

Herein, we report the creation of a new IPN of NorHA and collagen, providing control over matrix degradability and stiffness, independent of total protein and polysaccharide density (**Figure 1**). We demonstrate the formation of a viable vascular network from iPSC-EPs in this platform, quantified by an improved open-source computational pipeline [34]. These data represent a significant advance toward engineered vascular tissue from a patient-specific cell type.

**Figure 1:**
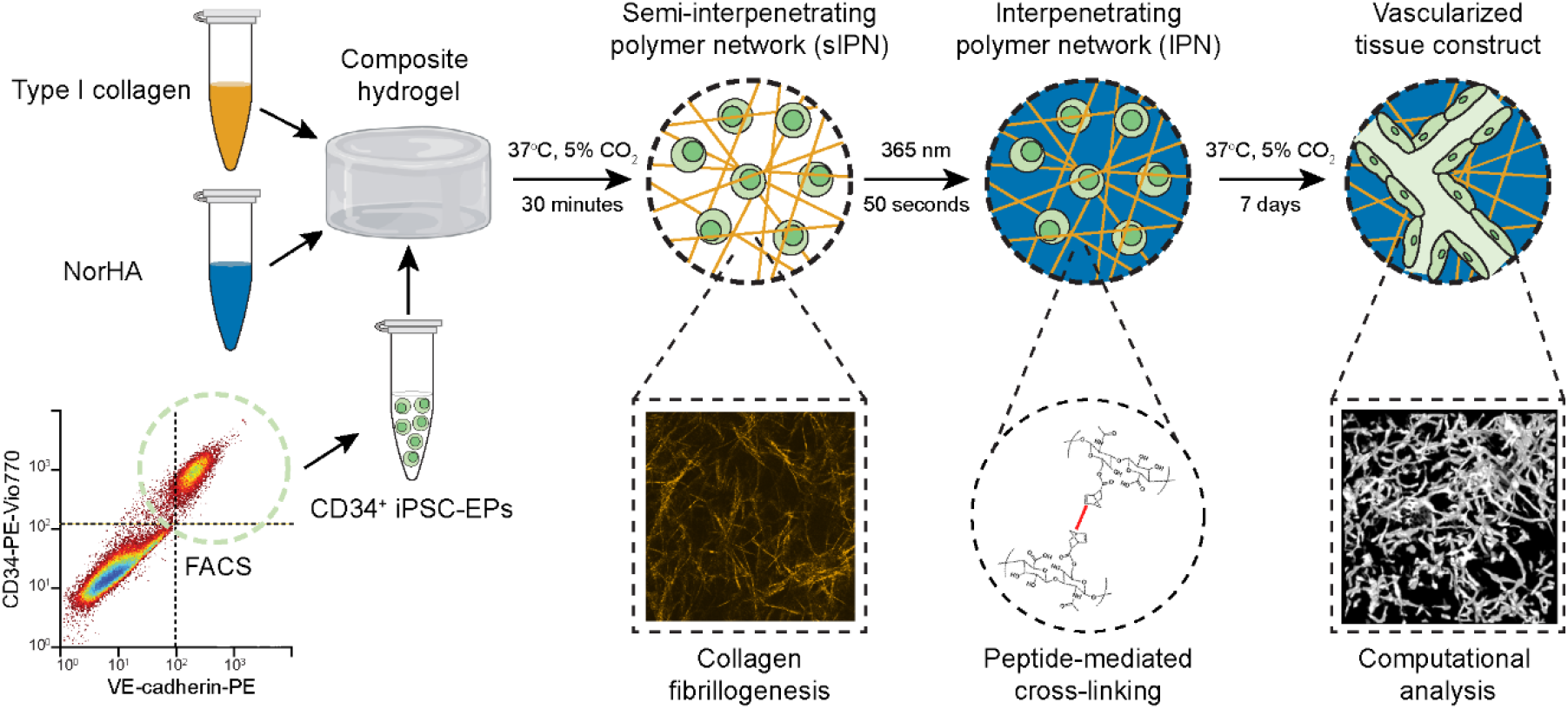
A schematic outlining the creation of UV-mediated collagen-NorHA-interpenetrating polymer network hydrogels. CD34^+^ iPSC-EPs are encapsulated in composite hydrogels containing NorHA and collagen. Collagen first physically cross-links upon neutralization and incubation at 37°C for 30 minutes. Norbornene-modified hyaluronic acid (NorHA) then cross-links in the presence of the photoinitiator LAP when exposed to UV for 50 seconds. In one week, encapsulated CD34^+^ iPSC-EPs form interconnected microvascular networks. These network topologies are imaged on a confocal microscope and robustly quantified by our novel, open-source computational pipeline. Composite hydrogel image was downloaded from BioRender.com.

## 2. MATERIALS AND METHODS

### 2.1 Functionalization of hyaluronic acid (HA) with norbornene (NorHA)

NorHA was synthesized following the procedure from Wade *et al.* with small modifications (**Figure S1A**) [35]. Briefly, HA was first converted to a tetrabutylammonium salt (HA-TBA). Sodium HA (Na-HA, 72 kDa, Lifecore) was dissolved in deionized (DI) water at 2 wt% with Dowex 50W ion exchange resin (3:1 resin to HA by weight) and stirred for 5 hours. The resin was filtered out, and the solution was titrated to pH 7.05 with tetrabutylammonium hydroxide, frozen, and lyophilized. Next, HA-TBA (2 wt%), 5-norbornene-2-carboxylic acid (12:1 to HA-TBA repeat unit), and 4-(dimethylamino)pyridine (3:1 to HA-TBA repeat unit) were dissolved in anhydrous dimethyl sulfoxide (DMSO) under N_2_ at 45°C. Next, di-tert-butyl dicarbonate (Boc_2_O, 1.6:1 to HA-TBA repeat unit) was added to the flask via syringe, and the reaction proceeded overnight. The reaction was then quenched with 4x excess DI water and dialyzed for three days. On day 3, 1 g NaCl per 100 mL was added, and the solution was precipitated into cold (4°C) acetone. The precipitate was spun down and re-dissolved into DI water, dialyzed for an additional week, frozen and lyophilized. Proton nuclear magnetic resonance spectroscopy (^1^H NMR) confirmed ~75% of HA repeat units had been functionalized with norbornene (**Figure S1B**).

### 2.2 Peptide synthesis and purification

RGD (GCGYGRGDSPG), an enzymatically degradable (DGD) peptide (KCGPQGIWGQCK), and a less degradable scrambled (SCR) peptide (KCGGIQQWGPCK) were synthesized using Rink Amide polystyrene resin (0.72 mmol/g from Gyros Protein Technologies) on a Prelude X automated peptide synthesizer (Gyros Protein Technologies). Traditional Fmoc-mediated coupling methods were used. Once complete, both syntheses were cleaved from the resin with 15 mL of 95:2.5:2.5 trifluoroacetic acid:water:triisopropylsilane for 4 hours. The peptides were dissolved at 10 mg/mL in 20:80 acetonitrile: water with 0.1% trifluoroacetic acid, and purified with a C18 column on a Dionex UltiMate 3000 UHPLC using a 25 min gradient of acetonitrile in water (20%-100%, 10 mL/min). After lyophilization, masses were verified via MALDI-TOF (Applied Biosystems - Voyager-DE™ PRO) (**Figure S2**).

### 2.3 Single-polymer and IPN hydrogel fabrication

Lyophilized NorHA was weighed in a 1.5 mL Eppendorf microcentrifuge tube and UV sterilized for 15 minutes. 20 mg/ml RGD, 40 mg/ml DGD, and 40 mg/ml SCR solutions were prepared by dissolving and vortexing lyophilized peptide in Endothelial Growth Medium 2 (EGM-2, PromoCell). The photo-initiator, lithium phenyl-2,4,6-trimethylbenzoylphosphinate (LAP), was dissolved in EGM-2 at a final concentration of 0.05 wt%. RGD was added to the lyophilized NorHA at a final concentration of 2 mM; DGD/SCR solution was added to achieve a final cross-linking ratio of 25/50/75/100% (of total norbornenes). The resulting solution was spun on a mini-centrifuge (Sprout, Fisher Scientific) for one hour and vortexed until transparent and homogenous.

Collagen hydrogels were fabricated as described previously [34,36]. 10X Medium 199 (ThermoFisher Scientific) and ice-cold 1 wt% collagen dissolved in acetic acid (Type 1, rat tail, Corning) were mixed and neutralized with 1 M sodium hydroxide (NaOH; Sigma-Aldrich), turning the solution bright pink. This solution was added directly to the NorHA solution containing RGD or DGD/SCR peptides. LAP was added to a final concentration of 0.05 wt%. 100 μL of the neutralized collagen-NorHA-cell suspension was then pipetted into circular silicone molds placed on Sigmacote-treated glass slides and allowed to solidify for 30 minutes at 37°C and 5% CO2. The hydrogels were then exposed to 365 nm light at an intensity of 10 mW/cm^2^ for 50 seconds (Omnicure Series 1500). The final hydrogels consisted of 1% wt NorHA and 0.25 wt% collagen.

### 2.4 Collagen fibrillogenesis and turbidity measurements

To quantify turbidity and thus gain a measure of the kinetics of type I collagen fibril assembly, hydrogels containing collagen were pipetted into a 96-well plate (Corning) and analyzed on a Cytation 3 Cell Imaging Multi-Mode Reader (BioTek Instruments). Absorbance readings at 313 nm [37] were taken every 30 seconds for a 30 minutes duration. To calculate the rate constants for the resulting kinetic growth curves, we solved and rearranged a simple first-order differential equation as described by Yan *et al.* to obtain the following linear equation [38]:

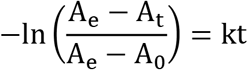

Where A_e_ is the absorbance at equilibrium, A_t_ is the absorbance at a given time t, and A_0_ is the absorbance at t = 0. By plotting the natural logarithmic expression against t, the rate constant can be estimated from a linear regression. To measure the final turbidity of the collagen or IPN hydrogel, samples analyzed on the Cytation 3 Imaging Multi-Mode Reader were exposed to 365 nm irradiation for 50 seconds at 10 mW/cm^2^ and then their absorbance was measured again at 313 nm.

### 2.5 Rheological measurements

Dynamic shear moduli and stress relaxation properties were measured on a TA Instruments HR-2 rheometer with an 8 mm parallel plate geometry. We first generated semi-IPNs (sIPN) by letting the collagen cross-link for 30 minutes at 37°C in a humidified environment in the presence of linear NorHA. The resulting sIPNs were placed on the rheometer quartz plate after 30 minutes at 37°C and 5% CO2 to ensure maximum contact. The NorHA was then gelled *in situ* upon exposure to UV light (365 nm irradiation for 50 seconds at 10 mW/cm^2^). A frequency sweep was performed from 0.1 to 100 Hz to determine the linear viscoelastic range; all subsequent modulus measurements were acquired at 1 Hz and 1% strain. To measure the relative stress relaxation properties of the collagen hydrogels, a 1% step strain was applied in shear, and the stress was recorded for 5 minutes.

### 2.6 Swelling ratio measurements

IPN hydrogels made *in situ* on the rheometer were subsequently transferred to a 12 well plate and allowed to swell in DI water at 37°C for 48 hours. Hydrogels were weighed in a 1.5 mL Eppendorf microcentrifuge tube (M_w_), frozen at −80°C, lyophilized (Labconco), and weighed again (M_D_). The degree of mass swelling (Q_m_) was calculated by calculating the ratio of the hydrogel wet mass (M_w_) to the hydrogel dry mass (M_D_):

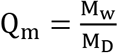

### 2.7 Confocal imaging of single-polymer and IPN hydrogels

After fabrication in an 8-well Nunc™ Lab-Tek™ II Chamber Slide™ System (ThermoFisher Scientific), single-polymer and IPN hydrogels were incubated with an HA binding protein derived from bovine nasal cartilage (EMD Millipore) as previously described by Suri & Schmidt [31]. Subsequently, the hydrogels were incubated with streptavidin (Cy5) overnight and washed extensively with PBS. The microarchitecture of single-polymer and IPN hydrogels was imaged using an inverted confocal microscope (Olympus FV3000) equipped with 20X, NA 0.75 objective. Collagen fibers were imaged under reflectance mode by illuminating with a 488 nm laser line and reflected light was collected by setting detector channel to acquire wavelengths between 485-495 nm. Confocal fluorescence images of Cy5-labeled hyaluronic acid were acquired with excitation wavelength at 640 nm and emission wavelength collected at 680−750 nm. All images were taken at a scan speed of 2 μs/pixel with a scan size of 1024 x 1024 pixels, and each line was averaged four times to produce the final images.

### 2.8 Generation and purification of CD34 expressing (CD34^+^) iPSC-EPs

iPSCs (DF19-19-9-11T) purchased from WiCell Technologies were maintained on vitronectin-coated plates in Essential 8 medium (E8, ThermoFisher Scientific), and passaged when the colonies reached ~75-80% confluence. iPSCs were differentiated into CD34^+^ iPSC-EPs following an established low-serum medium protocol [39]. Five days after CHIR induction (LC Technologies), cells were purified by methods we have described previously [34]. Briefly, the resulting confluent monolayer of differentiated cells was dispersed into a single-cell suspension with Accutase, tagged with a CD34-PE antibody (Miltenyi Biotec), and purified on a fluorescence-activated cell sorting (FACS) instrument (S3e Cell Sorter, Bio-Rad).

### 2.9 Vascular network visualization via confocal fluorescence

The microvascular networks in single-polymer and IPN hydrogels were visualized by staining the fixed cells with rhodamine-phalloidin, using previously established techniques [40]. Briefly, the hydrogels were fixed in a 4% paraformaldehyde and 5% sucrose solution for 10 minutes, washed three times with phosphate-buffered saline (PBS) supplemented with 300 mM glycine, and then permeabilized with 0.5% Triton-X for 10 minutes. After three washes with PBS supplemented with 300 mM glycine, the fixed hydrogels were incubated in PBS with 1% bovine serum albumin and 0.5% Tween-20 for 30 minutes and then with 1:40 rhodamine-phalloidin (ThermoFisher Scientific) overnight. After extensive washing with PBS, the hydrogels were transferred to an μ-Slide Angiogenesis 15-well imaging slide (Ibidi). The microvascular networks were imaged on a confocal microscope (Olympus FV3000) having a 10X, NA 0.3 objective. Fluorescence micrographs of microvessels stained with rhodamine-phalloidin were acquired with an excitation wavelength at 561 nm and emission wavelength at 580−650 nm. Z-stacks were acquired at 25 μm intervals, and at least three ROIs (~1.62 mm^2^ cross-sections) across the full depth of the hydrogel were acquired per technical replicate; at least three technical replicates were collected per experimental group.

### 2.10 Image segmentation and analysis of microvascular network topology

To analyze the microvascular networks’ topology (i.e., length and connectivity), we improved a computational pipeline that we had previously developed in ImageJ and MATLAB [34]. Briefly, confocal z-stacks that contained a cobblestone morphology on the surface of the gel were manually removed. Images were converted to an 8-bit scale, contrast compensated [41], binarized [42], and despeckled. The ImageJ “Analyze Particles” plugin removed particles with an area of fewer than 50 pixels and a circularity greater than 0.8, as these parameters tend to be indicative of dead cells. The filtered, binarized images were skeletonized and analyzed with a MATLAB algorithm that has been previously applied to analyze the quality of osteocyte networks in bone tissue [43] and to analyze the network topology of human umbilical vein endothelial cells (HUVECs) embedded in collagen hydrogels [44]. Briefly, each binary z-stack was skeletonized via an iterative algorithm and then converted into a network consisting of nodes and branches. In the previous version of this computational pipeline, we quantified the connectivity, vessel diameter, and the total number of branch points and vessels. With our improvements, we were able to quantify tortuosity, volume fraction, and the average diffusivity, i.e., the average distance from a vessel to a cell, from the skeleton and nodal topology. Each z-stack (~10-15 MB) required ~15 minutes to process on a Lenovo X250 ThinkPad equipped with 8 GB of RAM and an Intel Core i5-5200 CPU.

### 2.11 Statistical analysis

Statistical analysis was performed using a Student’s t-test (2 experimental conditions) or a one-way ANOVA (3 or more experimental conditions) followed by a Tukey’s multiple comparison test in Prism 6 (GraphPad). Significance is denoted as follows: * = p < 0.05, ** = p < 0.01, *** = p < 0.005, and **** = p < 0.001. Data are presented as mean ± standard deviation of at least two biological replicates unless indicated otherwise.

## 3. RESULTS

### 3.1 Rheological characterization reveals a two-step gelation mechanism and the decoupling of stiffness from bulk density

Initial experiments were conducted to confirm the independent cross-linking of collagen and NorHA, as any cross-reactivity between the two constituent polymers would indicate grafting onto a single polymer network, and not the formation of an IPN [45]. Type I collagen, lyophilized NorHA, and the degradable peptide cross-linker were reconstituted at a final concentration of 2.5 mg/mL, 10 mg/ml, and 2 mM, respectively, pipetted onto a TA Instruments HR-2 rheometer and then exposed to 10 mW/cm^2^ 365 nm irradiation for 50 seconds. The storage modulus of the hydrogels was measured across a range of frequencies to ensure that all subsequent measurements were acquired in the plateau range (**Figure S3**). NorHA and collagen, in the absence of the cross-linker, did not exhibit a storage modulus larger than 1 Pa, i.e., remained fluid for the duration of the experiment (**Figure 2A**). Interestingly, the addition of collagen to NorHA in the presence of a peptide cross-linker increased the final storage modulus of the sIPN to 50 Pa, which was significantly higher than that of 10 mg/mL NorHA and cross-linker alone (10 Pa). These experiments were conducted without NaOH, and therefore, the collagen did not form fibrils or physically cross-link in the acidic environment. Consequently, we sought to test if NorHA could be cross-linked after collagen monomers had assembled into polymeric fibrils and formed a physically cross-linked hydrogel. 2.5 mg/mL collagen hydrogels containing 10 mg/mL NorHA, 0.05 wt% photoinitiator (LAP), and varying amounts of cross-linking peptide were incubated for 30 minutes at 37°C. Afterward, these composite hydrogels were exposed to 10 mW/cm^2^ 365 nm irradiation for 50 seconds, and subsequently, the storage modulus was measured at 10-second intervals. We confirmed that collagen exhibited significantly larger storage than loss moduli, i.e., formed a physical hydrogel, in the presence of NorHA, and the storage modulus rose by up to an order of magnitude during UV exposure and remained stable afterward (**Figure 2B**). The final storage modulus of the IPN hydrogel increased monotonically in response to increasing cross-linker concentration (**Figure 3A**). Additionally, we substituted the degradable peptide cross-linker for a cross-linker that consisted of a scrambled amino acid sequence to reduce NorHA’s susceptibility to enzymatic degradation. When we varied the concentration of the scrambled cross-linker, the collagen hydrogel was still physically cross-linked, and the storage modulus of the final IPN also increased linearly with peptide concentration (**Figure 3A**). Finally, though the IPNs were considerably more elastic than single-polymer collagen hydrogels, they still dissipated roughly half of an applied load after 100 seconds, implying that the IPNs are viscoelastic (**Figure S4**). These rheological properties demonstrate that type I collagen and NorHA form IPN hydrogels via a two-step gelation process and the stiffness of the final IPN hydrogel can be varied simply by increasing the thiol:ene ratio of the crosslinking peptide, i.e., the stiffness of the hydrogel can be decoupled from the bulk polymer density of the final construct.

**Figure 2:**
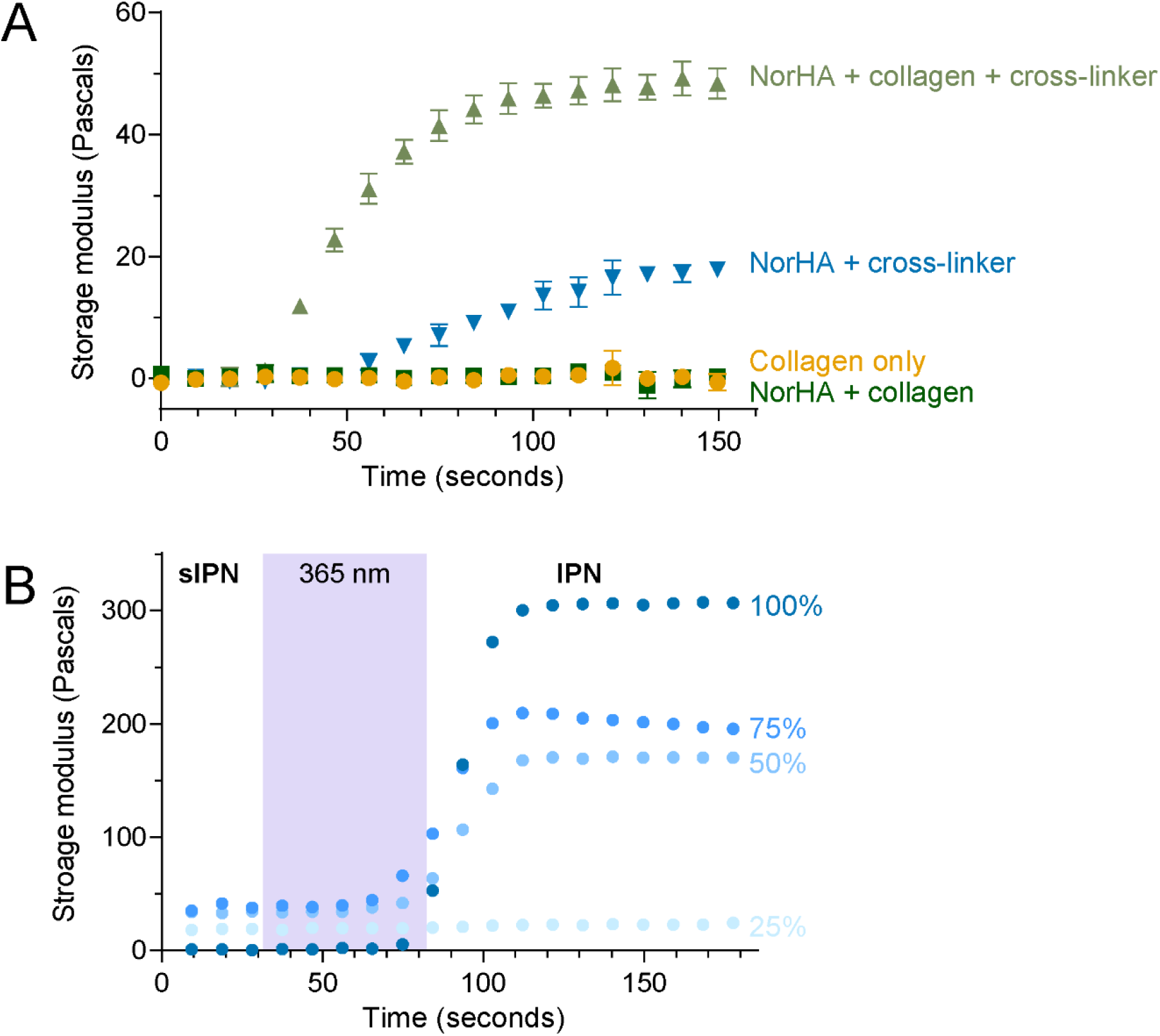
Rheological characterization of single-polymer and IPN hydrogels **(A)** Samples with non-neutralized collagen and NorHA were exposed to UV light, and the resulting storage modulus was measured via oscillatory rheology. **(B)** Composite hydrogels were synthesized; the collagen physically cross-linked for 30 minutes at 37°C. The resulting sIPN was then exposed to 365 nm radiation for 50 seconds, and the storage modulus was measured over the next 200 seconds.

**Figure 3:**
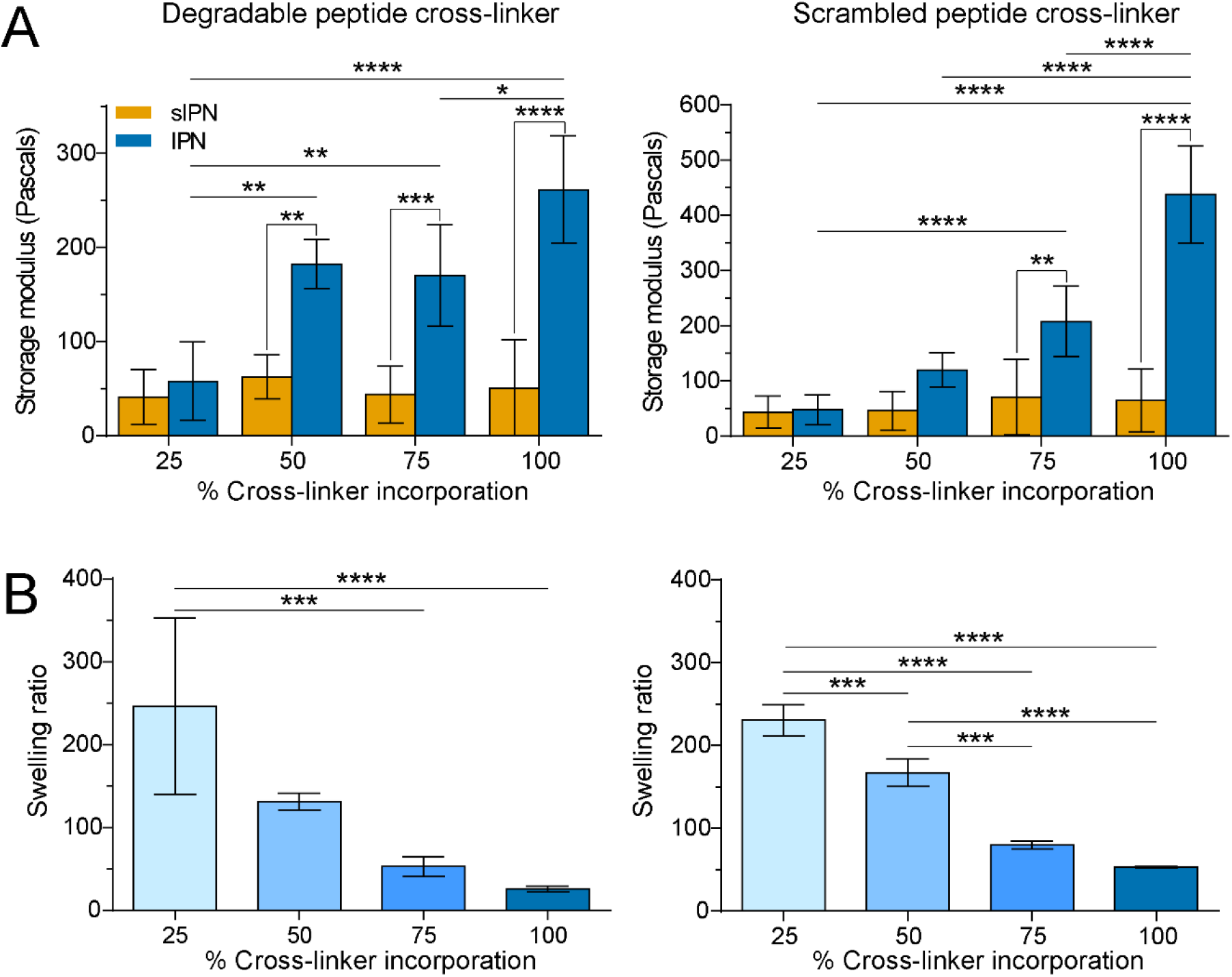
Characterization of IPN hydrogel stiffness and degradability **(A)** The storage modulus of composite hydrogels with differential cross-linking was measured via oscillatory rheology before exposure to UV (sIPNs, orange) and after UV (IPN, blue). **(B)** IPN hydrogels were swelled in PBS for 24 hours, weighed, lyophilized, and weighed again. From these two measurements, swelling ratios were calculated for differentially cross-linked IPN hydrogels with degradable and scrambled peptide cross-linkers.

### 3.2 The swelling ratio of IPN hydrogels decreases with increasing NorHA cross-linking

The swelling ratio quantifies the concentration of water remaining in the hydrogel at equilibrium. Critically, this parameter can give additional insights into the hydrophobicity of the hydrogel and can be used as an estimate of pore size and the relative diffusivity of the network structure [46]. After rheological testing, the hydrogels were swollen in water for 48 hours, weighed, frozen, and lyophilized for 48 hours. The swelling ratio was calculated from the ratio of the wet mass of the polymer to the dry mass, with a larger ratio indicating an increased degree of swelling. The swelling ratio decreased with increasing cross-linker concentration in the degradable hydrogels, ultimately decreasing fivefold when compared to collagen hydrogels (**Figure 3B**). When the degradable peptide was substituted with a significantly less degradable (“scrambled”) cross-linking peptide, the trend was nearly identical (**Figure 3B**). Therefore, we observed that increasing NorHA cross-linking decreased the swelling ratio, and the trend observed remained independent of the species of cross-linker used in these bioactive IPN hydrogels.

### 3.3 Fibrillogenesis in IPN hydrogels is slowed by the presence of HA and results in larger diameter fibers in increasingly opaque hydrogels

When exposed to elevated temperature and pH, type I rat tail collagen undergoes spontaneous fibrillogenesis and forms weak, physically cross-linked hydrogels. The resulting fibrillar microarchitecture is reminiscent of ECM fibrils, and the length, thickness, and orientation of collagen fibers impact the topology of vascular networks [47,48]. Though previous hydrogels consisting of HA and collagen have been reported, few studies have thoroughly analyzed the collagen network architecture in the resulting IPN. To analyze the kinetics of fibrillogenesis of the IPN hydrogels, the 313 nm absorbance of a neutralized collagen (2.5 mg/mL)/ NorHA (10 mg/mL)/ peptide cross-linker was measured at 30-second intervals at 37°C on an imaging plate reader. Notably, adding NorHA to collagen extends the fibrillogenesis lag phase by about 5-10 minutes (**Figure 4A**). To quantitatively compare the fibrillogenesis growth phases, we calculated the rate constant as described in the Materials and Methods and compared the values across all four conditions (**Table 1**) and found that the rate constants (~0.3-0.5 min^−1^) were similar across all conditions. The same hydrogels were then exposed to 10 mW/cm^2^ 365 nm irradiation for 50 seconds to measure their turbidity upon NorHA cross-linking. Strikingly, the hydrogels were noticeably more turbid when more cross-linker was incorporated into the hydrogel, and the absorbance values at 313 nm confirmed this trend (**Figure 4B**). Confocal reflectance was used to visualize the collagen network architecture of the IPNs. The addition of NorHA notably changed the network architecture; the collagen fibrils were less dispersed and appeared thicker than hydrogels composed of only collagen (**Figure 4C**).

**Table 1:** Estimated rate constants of fibrillogenesis for angiogenic hydrogels of differential composition.

| Hydrogel | Rate constant (1/minute) | R<sup>2</sup> |
| --- | --- | --- |
| Collagen | [-0.4453, -0.3618] | 0.83 |
| 25% NorHA-Collagen IPN | [-0.3247, -0.2871] | 0.9392 |
| 50% NorHA-Collagen IPN | [-0.4921, -0.4421] | 0.9461 |
| 75% NorHA-Collagen IPN | [-0.4559, -0.4058] | 0.9129 |
| 100% NorHA-Collagen IPN | [-0.2940, -0.2727] | 0.9657 |

**Figure 4:**
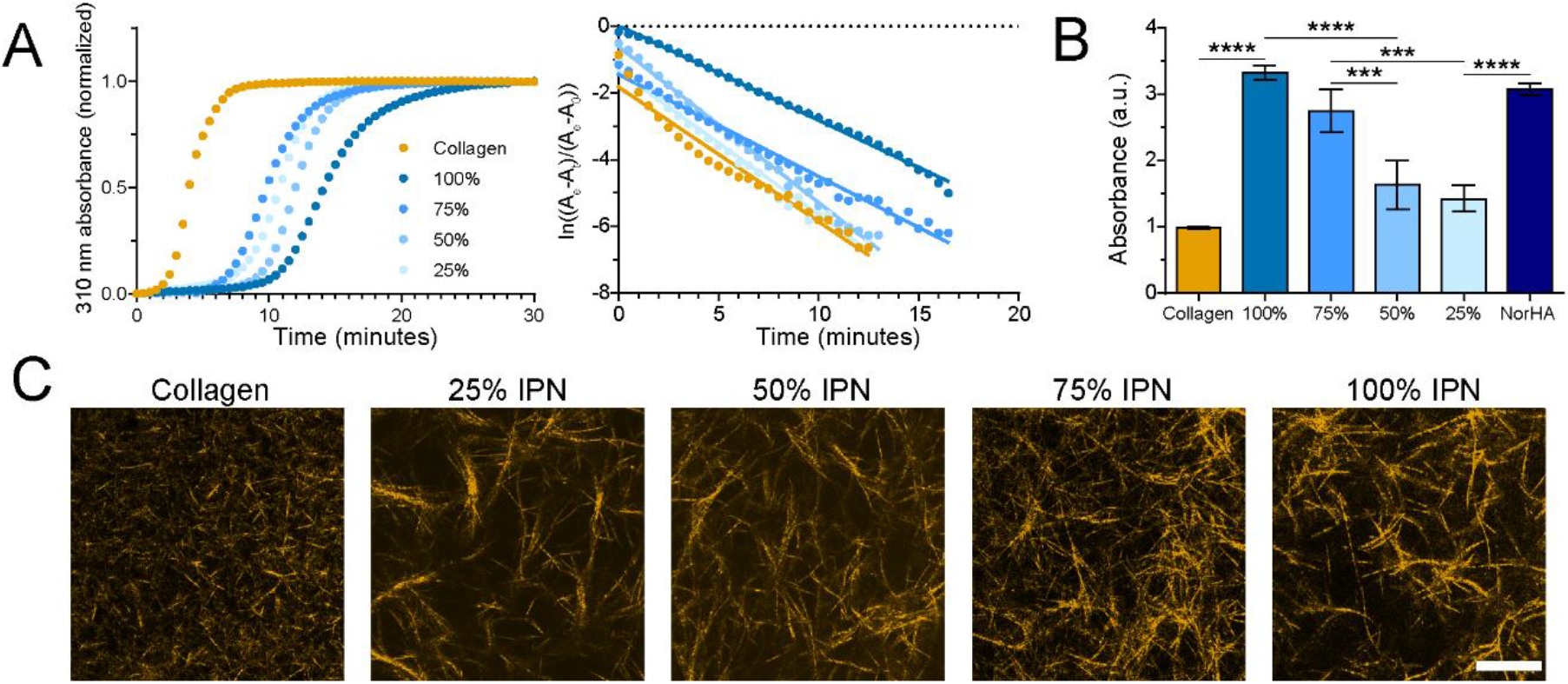
IPN fibrillogenesis kinetics and opacity varies with the cross-linking ratio. **(A)** Single-polymer and IPN hydrogels were prepared as previously described and pipetted into a 96-well plate. The absorbance was measured at 310 nm at 30-second intervals for 30 minutes. The rate constants of fibrillogenesis were extracted from and directly compared to evaluate if the kinetics differed between hydrogel formulations. **(B)** Immediately after, the same hydrogels were exposed to UV light at 10 mW/cm2 for 50 seconds. The absorbance was once again measured at 310 nm in triplicate and compared across conditions. **(C)** Collagen fiber architecture was imaged via confocal reflectance microscopy. The scale bar is 12.5 μm.

### 3.4 Optimization of cell-binding motifs and photoinitiator concentration to ensure cell viability and proliferation

As we have previously shown, CD34^+^ iPSC-EPs undergo apoptosis after encapsulation if Y-27632 (a ROCK pathway inhibitor) and VEGF are not immediately added to the culture medium [18]. Previously, we relied upon the native binding motifs found in collagen to encourage cell attachment and proliferation. To encourage the same phenomenon in collagen-NorHA IPN hydrogels, we grafted RGD, YIGSR, DGEA, or P15 to the backbone of 2D NorHA hydrogels to evaluate CD34^+^ iPSC-EP spreading and proliferation. We found that the cells remained viable and proliferated in all conditions, though the cells formed cobblestone-like colonies instead of lumenized structures (**Figure S5A**). RGD and YIGSR both encouraged significantly more proliferation than P15 and DGEA (**Figure S5B**). We, therefore, selected RGD as the cell-binding motif for future experiments.

To initiate NorHA cross-linking, we deployed LAP, a well-studied photoinitiator that has been previously demonstrated to be cytocompatible [49]. However, we found that the interaction of 0.5 wt% LAP and 365 nm irradiation resulted in considerable CD34^+^ iPSC-EP cytotoxicity (**Figure S6**). Considering these findings, we lowered the LAP concentration of 0.05 wt%; this concentration was used for all cell and material studies contained herein.

### 3.5 Lumenized microvascular networks develop from CD34^+^ iPSC-EPs in IPN hydrogels

CD34^+^ iPSC-EPs were sorted via FACS, encapsulated in NorHA-collagen IPN hydrogels, and cultured for one week in EGM-2 medium supplemented with 50 ng/mL VEGF, with daily medium changes. Interconnected lumenized microvascular networks formed after one week of culture. The networks were then fixed, stained with rhodamine-phalloidin, imaged on a laser-scanning confocal microscope, and analyzed with a custom computational pipeline in ImageJ and MATLAB to determine vessel diameter, number of branch points, diffusivity, tortuosity, volume fraction, and connectivity (**Figure 5**). We compared the vasculogenic potential of CD34^+^ iPSC-EPs in IPNs crosslinked with a degradable peptide (degradable IPNs), IPNs crosslinked with a scrambled peptide sequence (scrambled IPNs) and collagen-only hydrogels (the control). Upon viewing the binarized and filtered 3D volumes, the cell density in the collagen-only hydrogels was considerably higher, and LAP and UV reduced cell viability. The volume fraction, or the total percentage of the hydrogel occupied by the vascular network, was approximately four times larger in the collagen hydrogels than the other degradable IPN hydrogels. The diffusivity, average diameter, and the tortuosity of all microvascular networks were not statistically distinguishable among the degradable IPN hydrogels (**Figure 6**). Notably, the collagen network was highly interconnected, while the other degradable IPN hydrogels displayed limited connectivity due to lower cell density. After one week, degradable IPN hydrogels were approximately twice as thick as their collagen counterparts, which suggests increased resistance to degradation and subsequent compaction.

**Figure 5:**
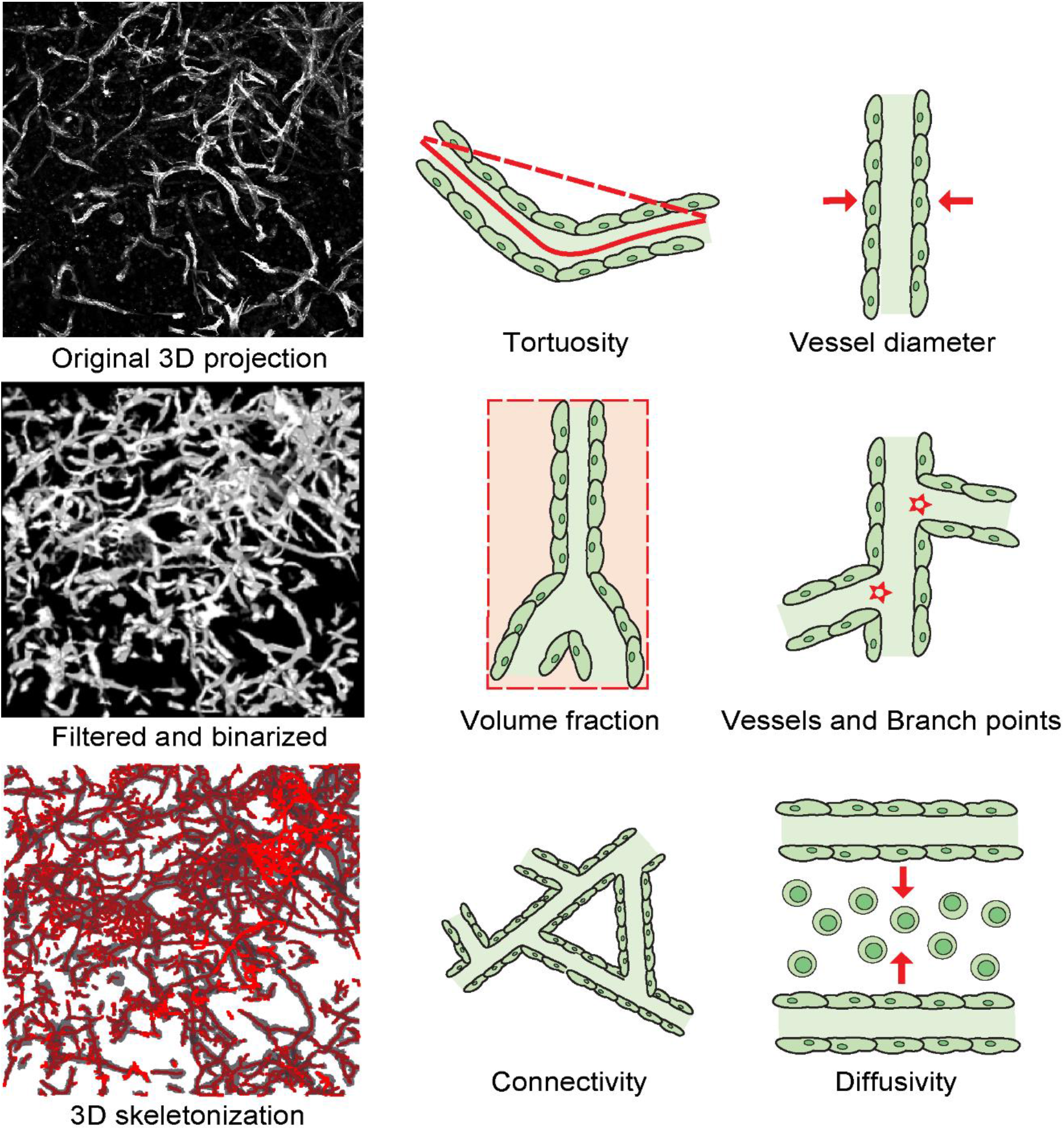
CD34^+^ iPSC-EPs were encapsulated in hydrogels at 2.2 million/mL and cultured for one week with daily medium changes. The hydrogels were then fixed, stained with rhodamine-phalloidin, and then imaged on a laser-scanning confocal microscope. The z-stacks were filtered, binarized, and converted into 3D representations using the “3D Viewer” plugin native to the ImageJ environment.

**Figure 6:**
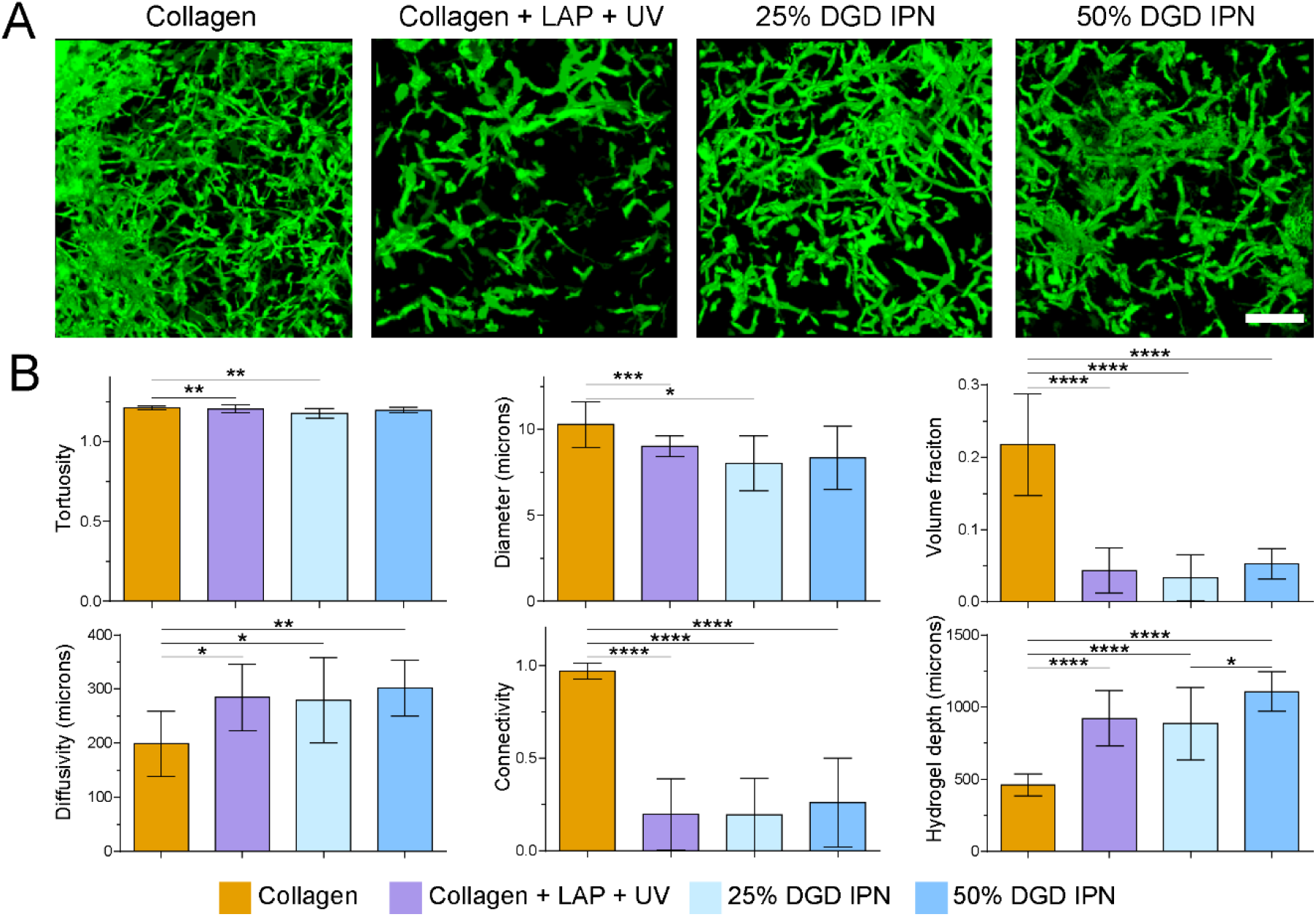
Robust iPSC-derived microvascular networks form in single-polymer and IPN hydrogels with degradable peptides **(A)** 3D representations of the final 3D, filtered, and binarized microvascular volumes were generated in ImageJ. The scale bar is 250 μm. **(B)** Network topology parameters were estimated with a custom MATLAB algorithm.

In contrast, the vascular parameters of microvascular vascular networks cultured in scrambled IPN hydrogels were markedly different. Collagen, 25%, and 50% scrambled IPN hydrogels occupied a similar volume fraction, were similarly diffuse, and exhibited similar connectivity (**Figure 7**). Furthermore, the 75% scrambled IPN hydrogels severely abrogated network growth, as most of the cells clumped and did not sprout into the surrounding collagen-hyaluronic acid matrix. Notably, while collagen generally contained more vessels and branch points than the degradable IPN hydrogels, the scrambled IPN hydrogels were markedly similar to the collagen samples. Specifically, the 50% SCR IPN contained a more significant number of vessels than the 25% SCR IPN, though this effect was reversed in the 75% SCR IPNs (**Figure 8**).

**Figure 7:**
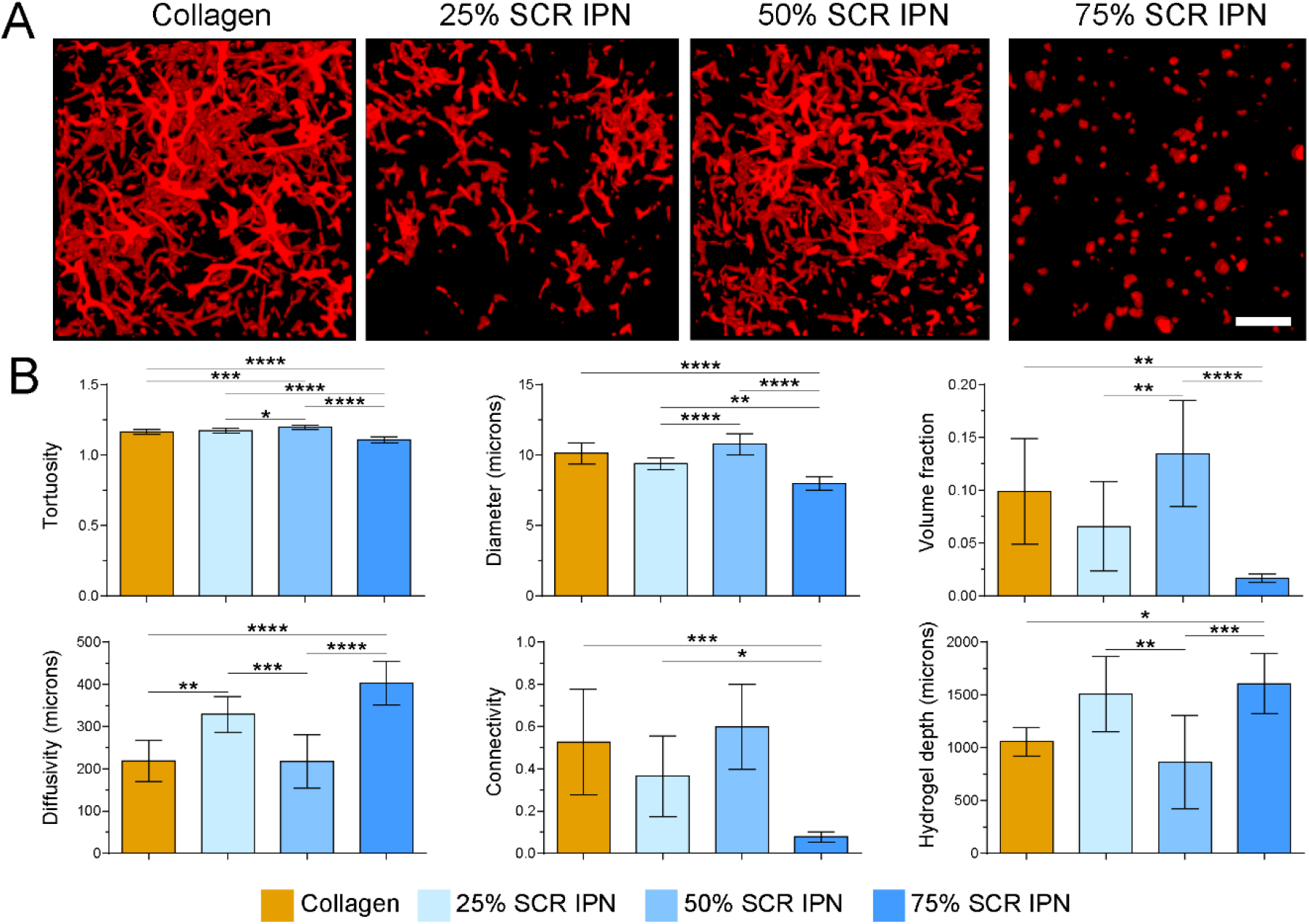
Robust iPSC-derived microvascular networks form in single-polymer and IPN hydrogels with scrambled peptides **(A)** 3D representations of the final 3D, filtered, and binarized microvascular volumes were generated in ImageJ. The scale bar is 250 μm. **(B)** Network topology parameters were estimated with a custom MATLAB algorithm.

**Figure 8:**
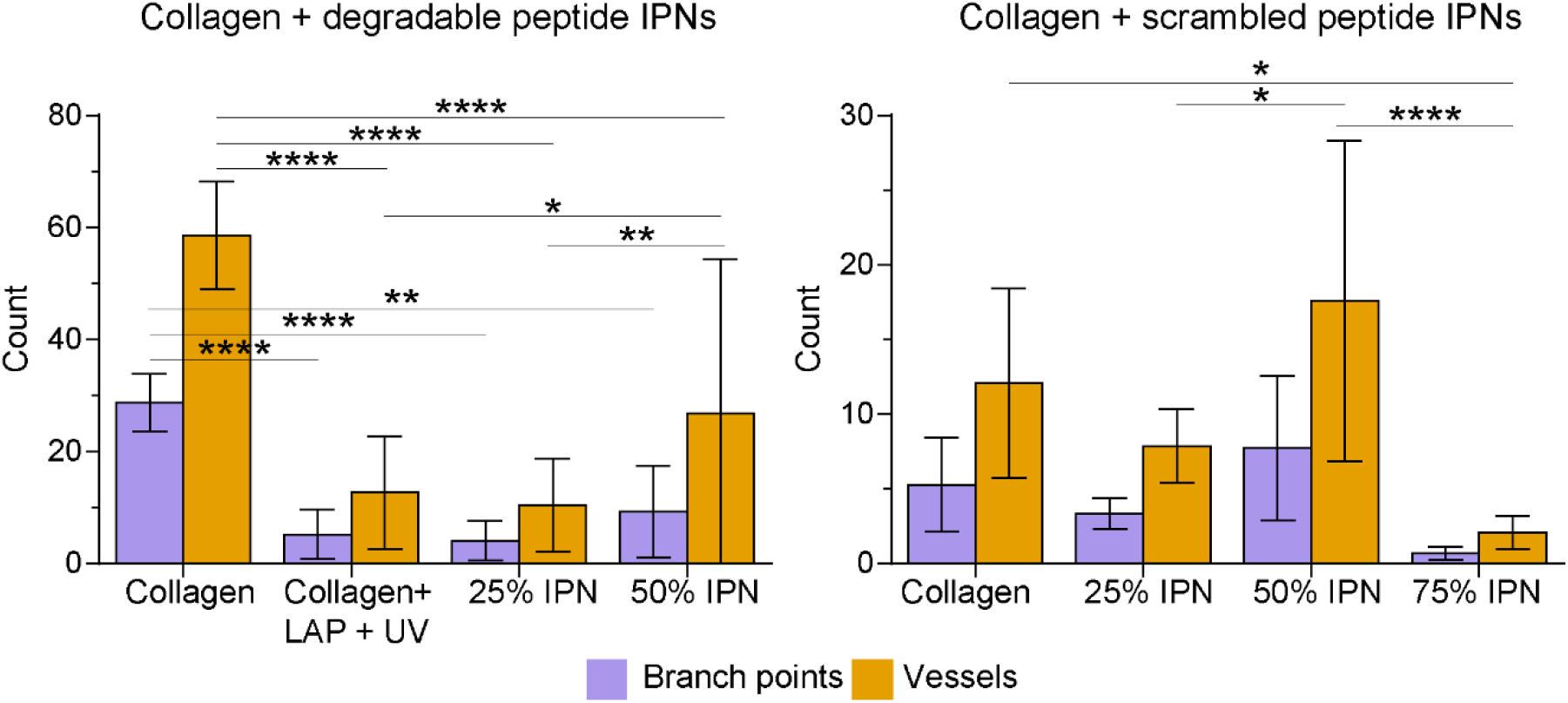
The branch points and links of the microvascular networks in single-polymer collagen hydrogels and IPNs containing degradable or scrambled peptide cross-linkers were extracted from the nodal topology generated by our MATLAB algorithm.

## 4. DISCUSSION

Previously, we had quantified iPSC-EP vasculogenesis in type I collagen hydrogels and confirmed that the network topology was dependent on the density of the surrounding matrix, which may have influenced the mean diffusion distance and degradability [18,34]. However, our experiments were limited to a single week due to severe collagen compaction. Furthermore, the stiffness of these collagen hydrogels remained well below physiological levels, and it proved difficult to modulate the stiffness of the construct without impacting other material parameters. We, therefore, sought to improve upon our previous work by synthesizing a novel collagen-based angiogenic biomaterial that would allow us to independently tune matrix stiffness and degradability while remaining stable in long-term culture.

Furthermore, it has been increasingly realized that many single-polymer biologically-derived hydrogels, such as fibrin or collagen, exhibit limited strength, and compact in long-term culture or under physiological conditions. To address these limitations, increasingly sophisticated hyaluronic acid-based hydrogels have been engineered to capture tissue properties such as viscoelasticity [50] and degradability [51]. These hydrogels have revealed extraordinary insights into cellular mechanotransduction and migration; however, these hydrogels often lack cell-scale topographical features, which are vital in guiding cell adhesion, migration, and proliferation [29]. Our system combines the precise cellular level control over stiffness and degradability that NorHA provides with the micro-fibrous and viscoelastic properties of collagen.

In our first experiment, we confirmed that we had synthesized a true IPN by showing that collagen does not cross-link with either NorHA or the peptide cross-linker. Also, we demonstrated that these IPN hydrogels were photo-responsive: upon exposure to 365 nm (UV) irradiation, the storage modulus of the composite hydrogels increased by approximately an order of magnitude. It is critical to note that the composite hydrogels gelled before UV exposure. This gelation implied that the collagen could self-assemble and thereby form a physically cross-linked network in the presence of NorHA. While others have detailed the synthesis of composite hydrogels composed of hyaluronic acid and collagen [30,32,33], we are the first, to our knowledge, to image and characterize the resulting collagen architecture. Notably, the collagen fibrils in the IPN hydrogels appeared larger and less disperse than fibrils in single-polymer collagen hydrogels. This architecture is reminiscent of cold-cast collagen, which indicates that this change in fiber formation could be accounted for by the additional time it takes the user to pipette and mix the seven constituent components of the IPN hydrogels at room temperature. It has been previously determined that cold-cast collagen, which is characterized by longer, thicker fibers, better promoted endothelial cell branching and collagen IV deposition [47].

Once we confirmed that our fibrous photo-responsive hydrogels formed an IPN via a two-step gelation process, we sought to vary physical and chemical parameters of the system systematically. We have noted in a previous review [5] that the impact of stiffness on engineered vasculature remains difficult to discern, though this confusion has often been rooted in a conflation of matrix density and matrix stiffness. Furthermore, stiffer substrates appear to stimulate tubulogenesis in 2-dimensions, but stiffer 3-dimensional scaffolds abrogate vascular network formation. Other groups have used novel techniques, from the glycation of collagen [20] to the creation of alginate-based IPNs [52,53], to combat this inherent interdependency. However, these systems are based on chemical modification of a bio-adhesive polymer, which may disrupt bio-functionality. Our system leverages collagen for bio-adhesive signaling and NorHA for mechanical support; furthermore, the bio-orthogonal cross-linking mechanism between norbornenes and small di-thiol peptides is user-defined, spatiotemporally controlled, and highly tunable. Thus, collagen-NorHA IPN hydrogels are uniquely poised to serve as a multimodal platform for probing the impact of stiffness on the vasculogenic potential of encapsulated iPSC-EPs. Specifically, we demonstrated control over IPN hydrogel stiffness, without changing bulk polymer concentration, or either NorHA or collagen.

Critically, we were also able to affect the degradability of the final IPN hydrogel by scrambling the sequence [27] of the otherwise degradable peptide. Increasing the concentration of the degradable and scrambled peptide led to an increase in storage modulus as well as an expected decrease in swelling ratio. In recent years, an increased emphasis has been placed on introducing viscous behavior into otherwise elastic gels, as stress dissipation is critical to cellular mechanotransduction. We found that increasing stiffness also increased the elastic nature of the IPN hydrogel. This increase in elasticity was expected, as the covalent thiol-ene linkages in the NorHA network impeded the ability of collagen to dissipate stress; however, some stress relaxation ability was preserved, and even the stiffest IPNs retained their viscoelasticity.

LAP is a widely deployed small molecule photoinitiator that induces gelation in synthetic and biological polymers while being considerably less toxic than its counterpart, I2959 [49]. However, at a concentration of 0.5 wt% [54], we observed that the interaction of 365 nm light and LAP proved cytotoxic to CD34^+^ iPSC-EPs. Interestingly, LAP and 365 nm light individually were cytocompatible, which leads to our speculation that free radicals produced by LAP are responsible for this observed cytotoxicity. When the concentration of LAP was reduced to 0.05 wt%, we observed considerably less cytotoxicity while still allowing the crosslinking reaction to occur. To further optimize cell viability, we tested different cell-binding peptide motifs that are traditionally contained in collagen helices [55]. Though collagen retains the motifs necessary for cell recognition and binding, we wanted to ensure that the entirety of the final IPN hydrogel was bioactive. RGD and YIGSR both increased CD34^+^ iPSC-EP binding and subsequent proliferation, and we chose RGD for use in all remaining studies.

After a week of culture, we found that cells in the 75% and 100% IPN hydrogels were not viable and did not form any visible structures. We speculate that this effect could be explained by a reduced diffusivity in the NorHA network, which would be compatible with the trend in swelling ratios we observed. In contrast, cells in the 25% and 50% IPN hydrogels with likely significantly larger mesh sizes, underwent tubulogenesis and formed lumenized structures that were stable for more than two weeks. Also, the IPN hydrogels were almost twice as thick at the end of the experiment, indicating that these composite hydrogels underwent considerably less compaction than single-polymer hydrogels. This difference in hydrogel thickness may be due to the increased stiffness of these IPNs which can better resist the forces exerted by the cells as they proliferate and migrate throughout the IPNs. However, it can also be due to decreased cell viability in UV-exposed hydrogels containing LAP.

In our previous work, we developed and deployed a computational pipeline based in ImageJ and MATLAB to analyze the topology of the resulting vascular networks. We made further improvements to our algorithm and were able to extract additional parameters: diffusivity, volume fraction, and tortuosity. For most conditions, the IPN hydrogels and the collagen-containing LAP that was exposed to UV encouraged the growth of microvascular networks with similar topologies. Though the collagen hydrogels appear to be more conducive to network formation (e.g., higher volume fraction and connectivity), this is partly due to their compaction, which compresses a similar number of vessels into a smaller total volume, making the nascent vascular network appear denser. This effect may also be partly due to the higher number of viable cells at the start of the experiment. Despite these effects, the network topology in the IPNs is approximately similar to that of the networks in collagen. Broadly, increasing the concentration of the peptide cross-linker resulted in an increased vasculogenic response until 75% of the cross-linking sites were occupied; at this occupancy, we observed severely reduced network formation. Notably, IPN hydrogels containing a scrambled peptide induced a more complex, interconnected vascular network topology than IPN hydrogels containing a more degradable peptide. We speculate that the increased stability of the scrambled IPN hydrogels allows for the cells to migrate, proliferate, and undergo tubulogenesis in a more stable microenvironment. The effect of relative degradability on iPSC-EP vascular network development is worthy of further elucidation.

## 5. CONCLUSION

We created a novel photosensitive IPN hydrogel that possesses the necessary material and bioactive properties to be a versatile platform for interrogating angiogenesis and vasculogenesis. EPs derived from iPSCs rapidly underwent tubulogenesis and formed interconnected vascular networks after one week in culture. The stiffness and degradability of the resulting IPN could be tuned by changing the species and concentration of peptide cross-linker. Furthermore, we demonstrated the power and utility of our improved computational pipeline that can robustly quantify network topology in three dimensions. Considering these findings, iPSC-EPs and angiogenic IPN hydrogels could serve as the biomimetic foundation of vascular models that may illuminate the mechanisms of cardiovascular pathologies and open the door for the development of new therapies.

## ACKNOWLEDGMENTS

We thank Shreya Ramesh and Katerina (Katya) Wittliff (Biomedical Engineering Department, The University of Texas at Austin) for their assistance with cell culture and experiments. We are grateful for illuminating discussions with Dr. Joshua Morgan (University of California, Riverside) on the computational analysis of 3D networks. We also acknowledge the use of shared research facilities supported in part by the Texas Materials Institute, the Center for Dynamics and Control of Materials: an NSF MRSEC (DMR-1720595), and the NSF National Nanotechnology Coordinated Infrastructure (ECCS-1542159).

---

**Figure S1:**
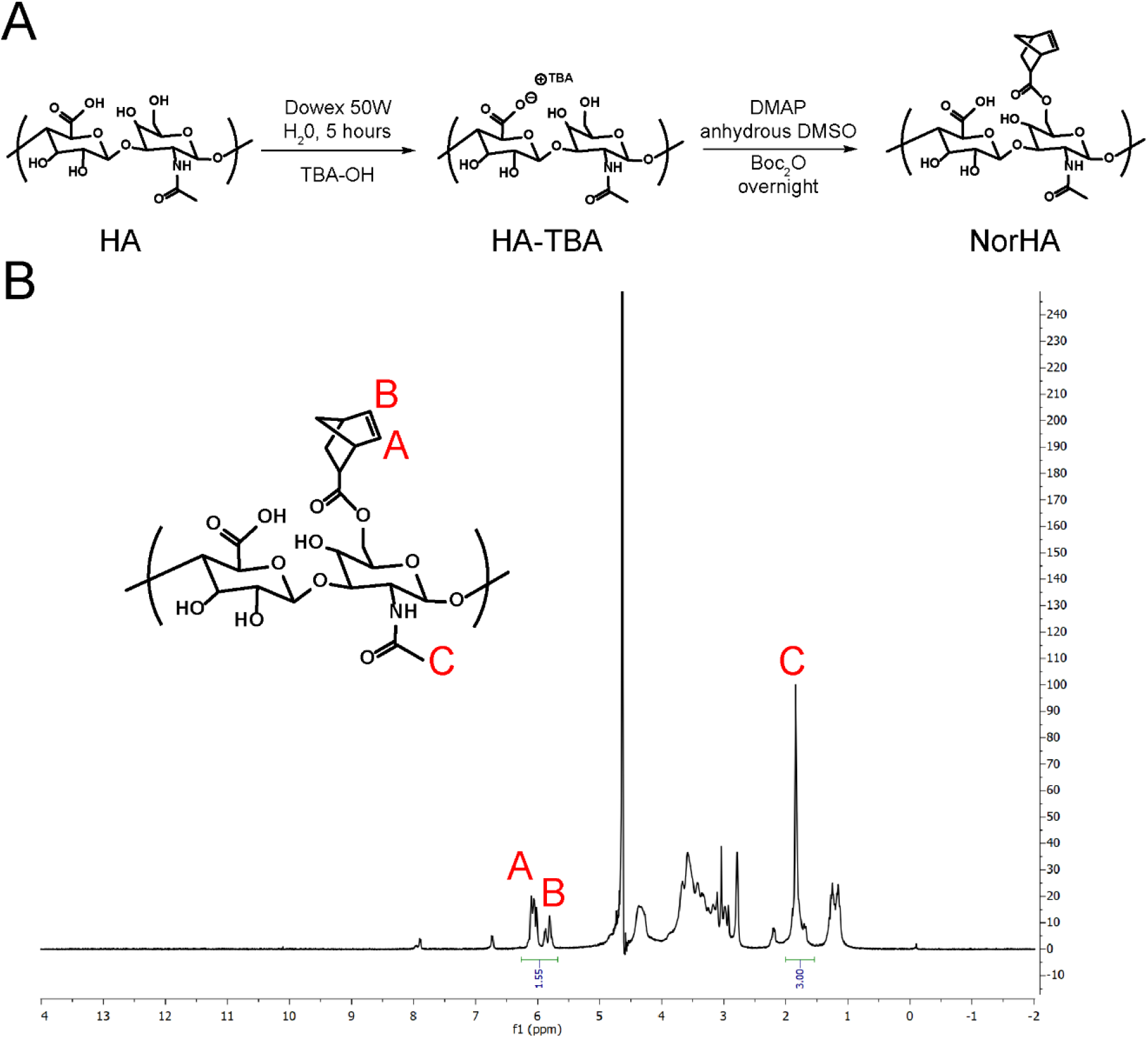
NorHA synthesis **(A)** Schema of the modification of HA to HA-TBA to NorHA. **(B)** ^1^H NMR spectrum of norbornene-modified hyaluronic acid (NorHA). The degree of modification was determined to be ~75%.

**Figures S2:**
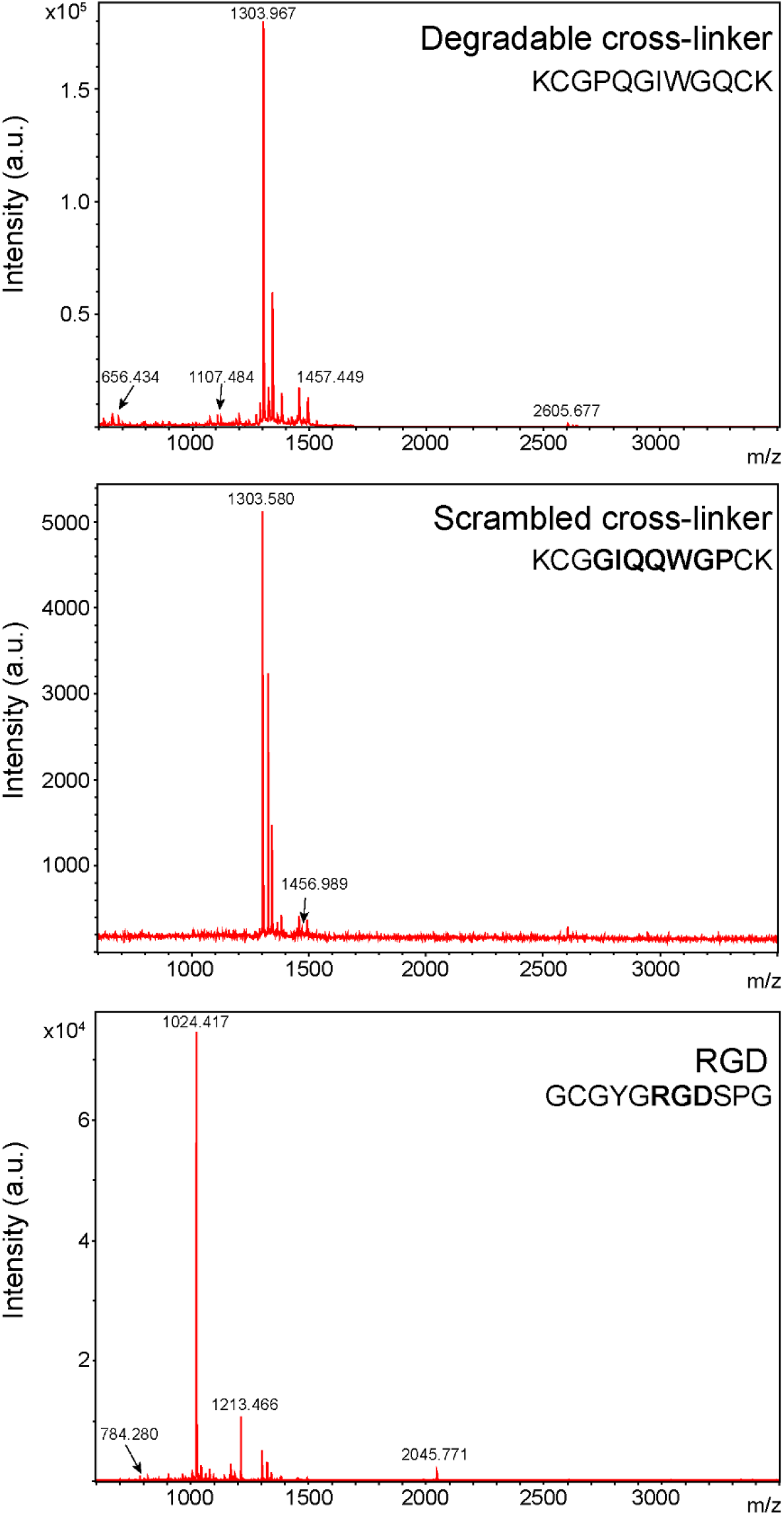
Molecular weights of the degradable/scrambled peptide cross-linker and RGD cell-binding motif were confirmed with MALDI.

**Figure S3:**
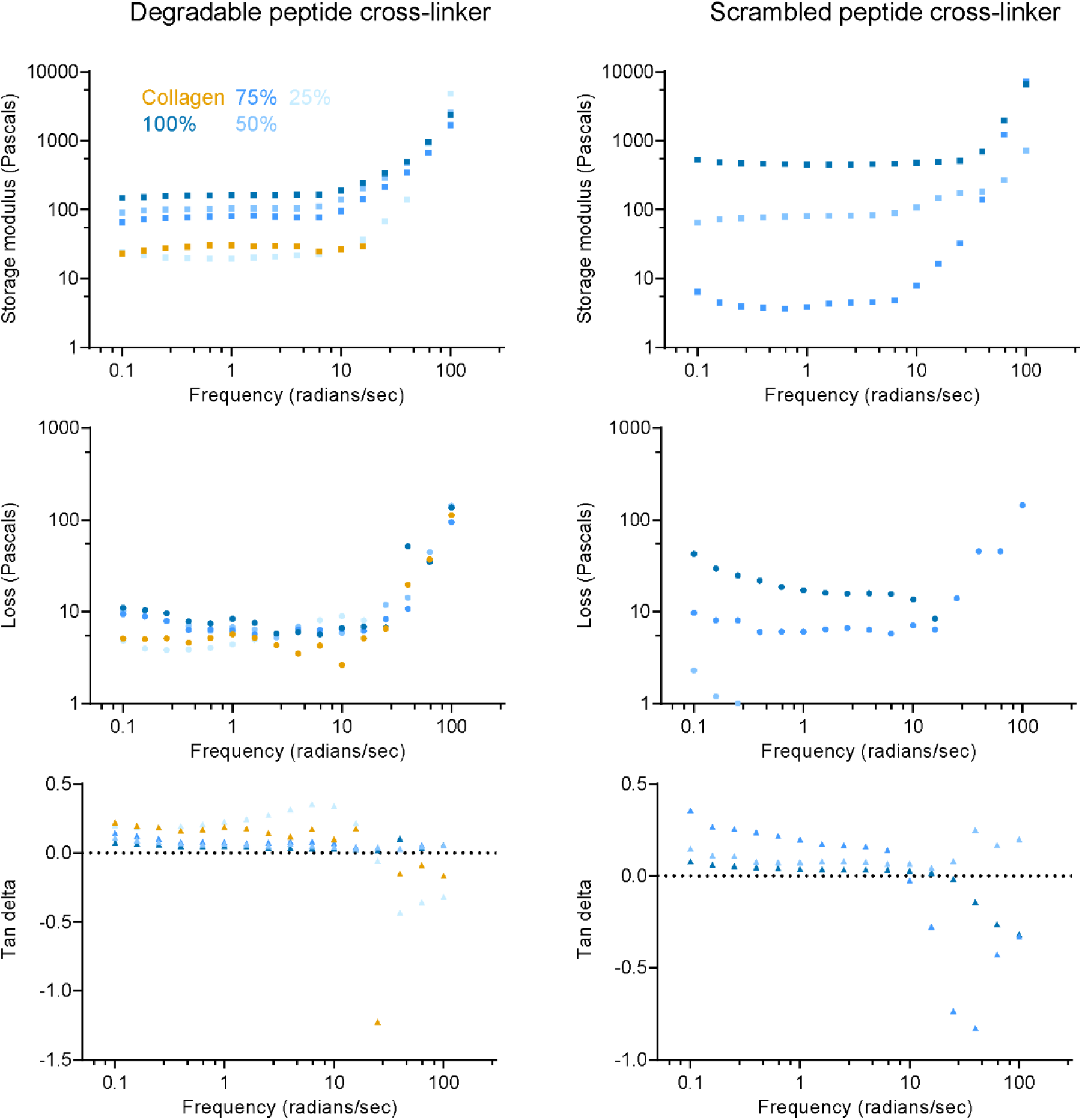
Storage moduli, loss moduli, and the tan delta of single-polymer collagen and IPN hydrogels plotted as a function of frequency.

**Figure S4:**
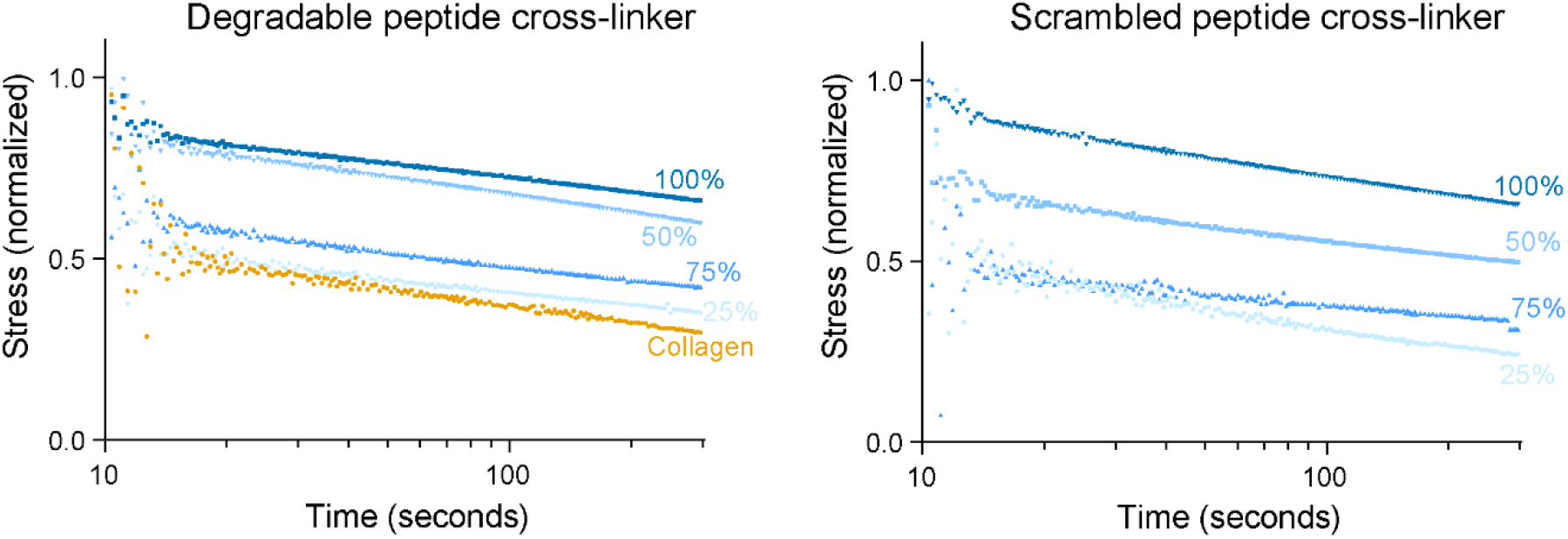
Stress relaxation tests conducted on single-polymer collagen hydrogels and IPNs with cross-linkers of degradable or scrambled sequences.

**Figure S5:**
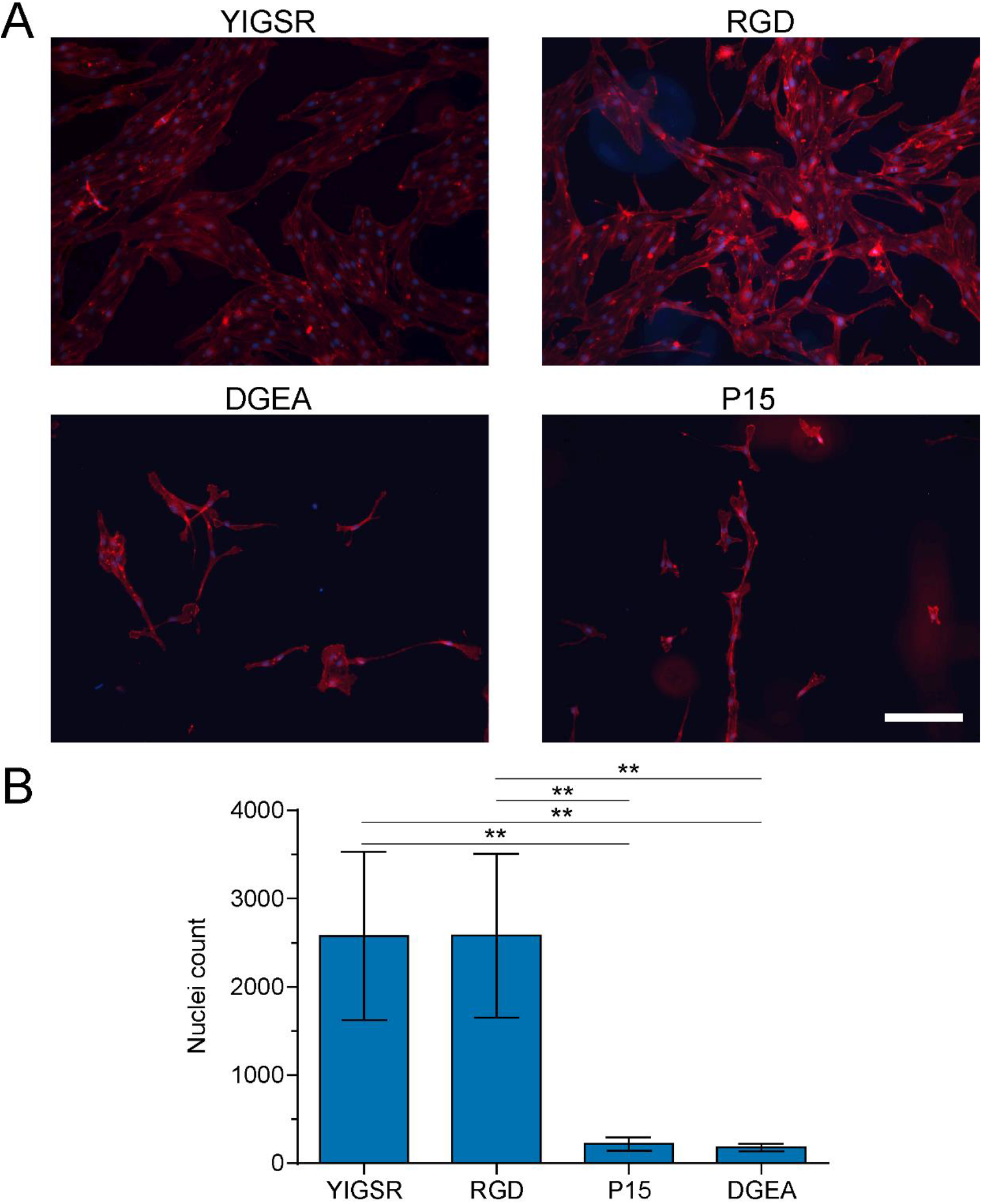
CD34^+^ iPSC-EPs preferentially attach and spread on YIGSR and RGD-modified NorHA substrates (A) After one week, cells were fixed and stained with rhodamine-phalloidin and DAPI to highlight the actin structure and nuclei, respectively. The scale bar is 200 μm. (B) The nuclei were imaged on a Cytation3 Imaging Plate Reader; the images were segmented, and the total number of nuclei were counted using the “Analyze Particles” Plugin native to ImageJ.

**Figure S6:**
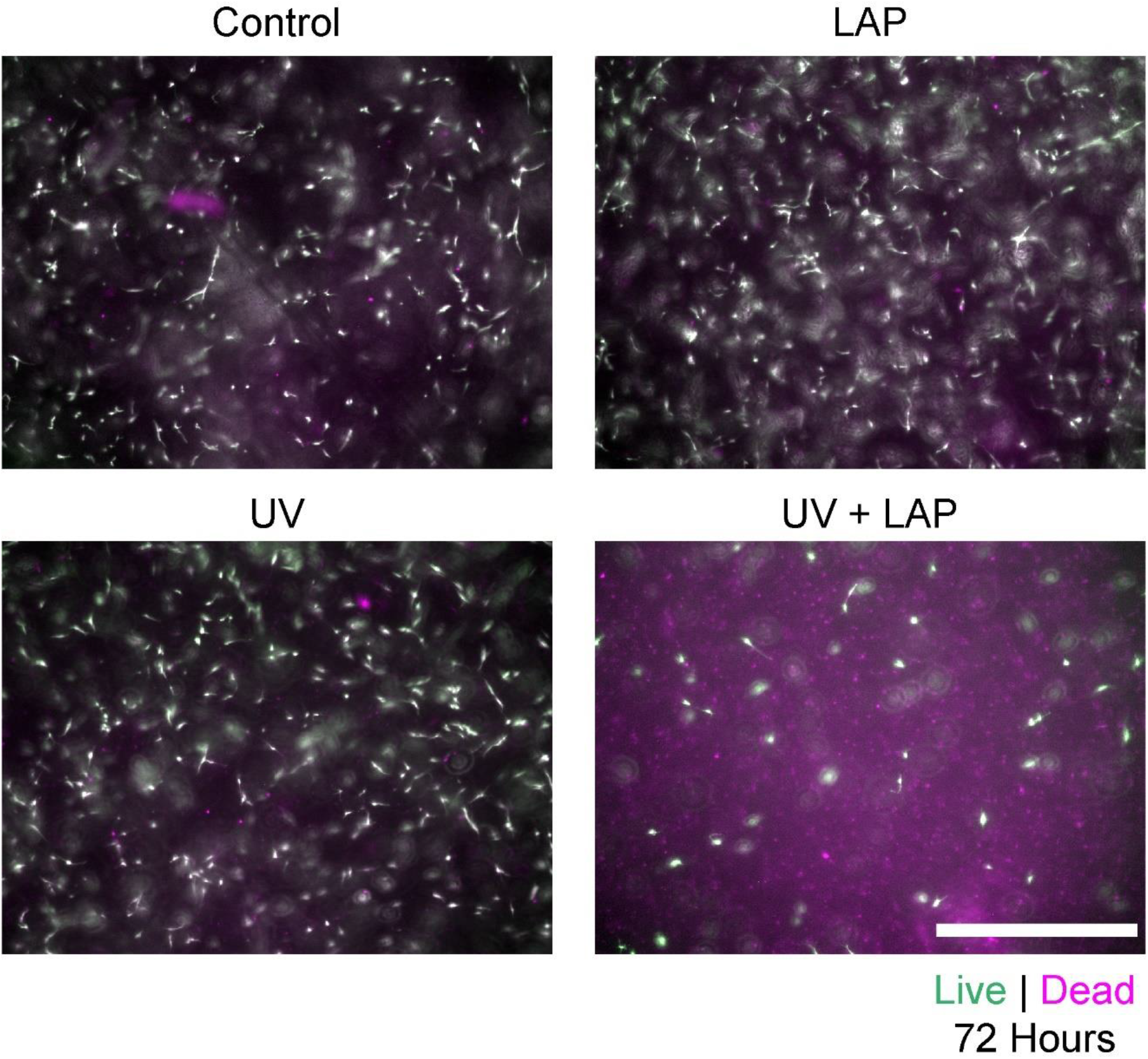
Interaction of 0.5 wt% LAP and 10 mW/cm^2^ UV induces CD34^+^ iPSC-EP apoptosis. Upon encapsulation in collagen hydrogels, cells were incubated standard culture medium, standard culture medium with LAP, exposed to UV, or exposed to UV in the presence of LAP. 72 hours after exposure, the cells were incubated with 2 mM calcein (green) and 2 mM ethidium-homodimer (magenta) to identify live and dead cells, respectively. The scalebar is 400 μm.

